# Understanding Admixture Fractions

**DOI:** 10.1101/008078

**Authors:** Mason Liang, Rasmus Nielsen

## Abstract

Estimation of admixture fractions has become one of the most commonly used computational tools in population genomics. However, there is remarkably little population genetic theory on their statistical properties. We develop theoretical results that can accurately predict means and variances of admixture proportions within a population using models with recombination and genetic drift. Based on established theory on measures of multilocus disequilibrium, we show that there is a set of recurrence relations that can be used to derive expectations for higher moments of the admixture fraction distribution. We obtain closed form solutions for some special cases. Using these results, we develop a method for estimating admixture parameters from estimated admixture proportion obtained from programs such as Structure or Admixture. We apply this method to HapMap data and find that the population history of African Americans, as expected, is not best explained by a single admixture event between people of European and African ancestry. A model of constant gene flow for the past 11 generations until 2 generations ago gives a better fit.

## Introduction

It is common in population genetic analyses to consider individuals as belonging fractionally to two or more discrete source populations. The proportion of an individual’s genome that belongs to a population is called that individual’s ‘admixture fraction’ or ‘admixture proportion’. Programs such as **Structure** (Pritchard et al., 2000), **Eigenstrat** (Price et al., 2006), **Frappe** (Tang et al., 2005), or **Admixture** (Alexander et al., 2009) can jointly estimate these admixture fractions for multiple individuals in a sample, along with the corresponding allele frequencies in each of the source populations. These admixture fractions are often presented in a ‘structure plot,’ an example of which is shown in Figure 1. We will henceforth refer to these methods as ‘structure analyses’. This approach has proven highly useful for understanding genetic relationships in many different species, e.g. humans (Rosenberg et al., 2002), cats (Menotti-Raymond et al., 2008), or pandas (Zhang et al., 2007). Other analyses reconstruct admixture tracts for each genome in the sample, by inferring the local ancestry of every position, or window, in each sampled genome (Tang et al., 2006; Maples et al., 2013). In this context, the admixture fraction for a genome is the fraction of its total length that is inherited from a particular source population.

**Figure 1.**
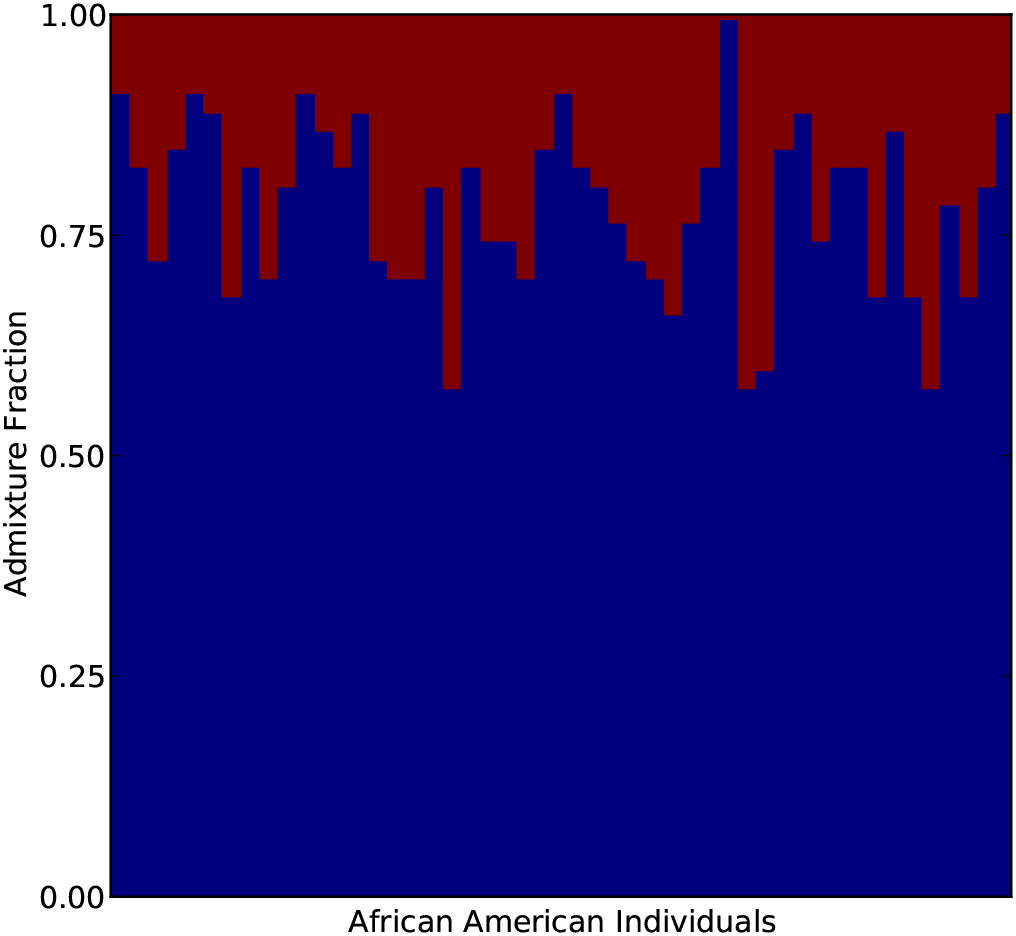
Admixture fractions for 49 African American individuals in the HapMap 3 data. Source population allele frequencies were estimated using 113 Yoruban and 111 European individuals.

Although structure analyses are not tied to any particular mechanistic model of population history and demography, the admixture fractions and admixture tracts are commonly interpreted to be the result of past admixture events in which modern populations were formed by admixture (or introgression) between ancestral source populations. The distribution of admixture tract lengths has been related to specific mechanistic models of admixture (Falush et al., 2003; Tang et al., 2006; Pool and Nielsen, 2009), and has been used to estimate times of admixture (Gravel, 2012). However, the admixture proportions themselves also contain information regarding admixture times. Following an admixture event, the variance in admixture proportions within a population will be high, but will there after decrease, and will eventually converge to zero in the limit of large genomes. The variance in admixture fractions among individuals contains substantial information about the time since admixture that can be used in addition to the tract length distribution. In some cases, this may be more robust than inferences based on tract lengths, because the length distribution of tracts is often difficult to infer, and is often not modeled accurately by the hidden Markov model (HMM) methods used to infer tract lengths (Liang and Nielsen, 2014). Even in cases where tract lengths can be accurately inferred, studies aimed at estimating admixture times should benefit from using both variance in admixture proportions among individuals and overall admixture tract lengths distributions.

Verdu and Rosenberg (2011) developed a method for computing moments of admixture proportions in a model in which admixed population is formed as a mixture between multiple source populations, allowing for arbitrary gene-flow from the source populations over a number of generations(*g*). They establish recursions for the moments of the admixture fractions and use these equations to determine how the mean and the variance changes through time in particular admixture scenarios. These moments are expectations for *single* individual’s admixture fraction and are averaged over the possibile genealogical histories of the population. As a result, they can be difficult to relate to data because replicates from multiple identical populations rarely are available. In this paper, we consider a different problem, the problem of calculating sample moments for admixture proportions obtained from individuals in one population.

We extend the model model in Verdu and Rosenberg (2011) to incorporate the effects of recombination and genetic drift by adding a a random union of zygotes component. Recombination is important because even if one half of a chromosome’s ancestors are from the first source population, it is unlikely that exactly one half of that chromosome’s genetic material is inherited from that population. Genetic drift is important because the individuals in a sample might share ancestors and, therefore, have more similar admixture fractions than expected by chance in a model without drift. The results developed in this paper should be directly applicable for quantifying the results of a structure analysis.

## The General Mechanistic Model

We start by considering admixture fractions in haploid genomes. These haploid admixture fractions can later be paired up to create diploid admixture fractions. The admixture fraction of a(haploid) genome *H_i_*, is the proportion of *H_i_* that is inherited from a particular source population. For notational simplicity, we only consider gene-flow only from one population into another. We will later discuss how to extend this model to multiple admixing source populations. We use the same mechanistic admixture model of Verdu and Rosenberg (2011), and will use its notation where possible. Finally, we use the random union of zygotes model, with a diploid population size of *N* (2*N* chromosomes), for genetic drift and recombination, and assume a sample size of *n* chromosomes from a single population.

In this model, a hybrid population of *N* diploid individuals forms in generation 1 from two previously isolated source populations. In this first generation, individuals in the hybrid population are from the first source population with probability *s*_0_ or from the second source population with probability 1 − *s*_0_. In generation *g* + 1, each chromosome is, independently, from the first source population with introgression probability s_*g*_, or from the hybrid population with probability 1 − *s_g_*. Chromosomes inherited from the hybrid population are the product of the recombination of the two chromosomes of one individual (zygote), chosen uniformly at random. Finally, these 2*N* chromosomes are paired up to form the *N* individuals in generation *g* + 1.

Finally, we let the stochastic process *A*(*ℓ*) represent the local ancestry along a chromosome as a function of *ℓ*, the physical position:

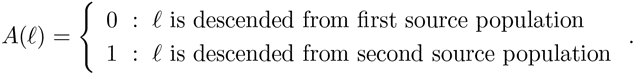

The fraction of the chromosome descended from the second source population is given by

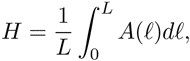

where *L* is the total length of the chromosome.

Assume that *g* generations after the start of admixture we have randomly sampled *n* chromosomes from the hybrid population and determined their corresponding admixture fractions, *H*_1(*g*)_, *H*_2(*g*)_, …, *H_n_*_(*g*)_. We are interested in the joint distribution of these *n* random variables. When *n* = 1 and as *L* → ∞, this is the admixture fraction considered by Verdu and Rosenberg (2011).

Because the *n* chromosomes have possibly overlapping geneologies, the admixture fractions are not independent. However, the joint distribution of the admixture fractions does not depend on their ordering, so they are exchangeable. As a result, they can be viewed as being identically and independently (*iid*) drawn from a random distribution 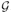. This random distribution can be interpreted as a function of the random genealogy of the entire hybrid population up to g generations in the past. When *g* is small, the genealogies of the *n* samples will be unlikely to differ from *n* nonoverlapping binary trees, so 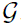 will be approximately constant. If *g* is large however, these genealogies are likely to overlap, and this will no longer be true.

Verdu and Rosenberg (2011) focus on moments of *H*_1(*g*)_, in particular on the mean and variance. However, because the admixture fractions are not independent, even as *n* → ∞, the sample mean and sample variance will converge to the mean and variance of 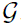, which are random quantities. For example,

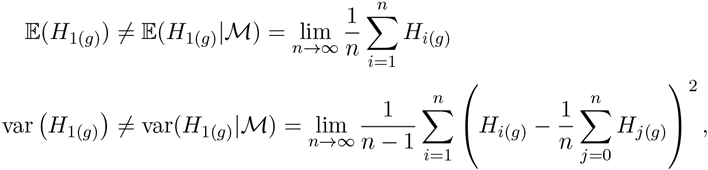

and similarly for higher-order moments. The moments of the admixture factions have two components: randomness from sampling the population genealogy, and randomness from the sampling of chromosomes. The expressions to left account for both, while the expressions to the right only account for the latter. Variances among individuals with in one population correspond to var(*H*_1(*g*)_|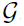), while variances over replicate populations correspond to var(*H*_1(*g*)_). This latter value will be larger than the expected sample variance calculated from multiple individuals sampled from the same population, and will rarely be useful for inference purposes.

In the following sections, we will show how the constants on the left-hand side, as well as expectations of the random variables on the right-hand side, can be derived for mechanistic models of introgression. By comparing these expectations to the observed admixture parameters from a sample, we will be able to construct a method of moments estimator for the parameters of the model.

Let *k*_1_ be the sample mean:

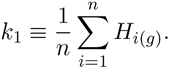

We can express its expectation in terms of the 1-point correlation function of *A*:

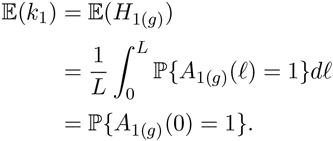

Similarly, let *k*_2_ be the unbiased estimator of the sample variance:

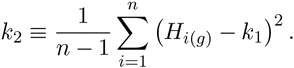

Its expectation is given by

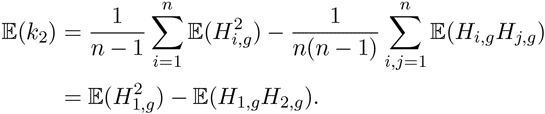

These expectations can be written in terms of two-point correlation functions of *A*:

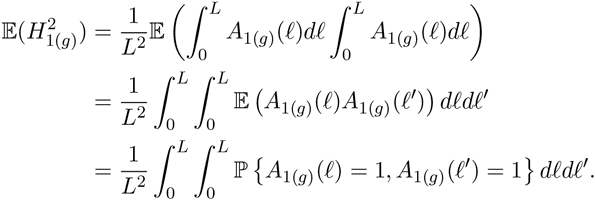

Similarly,

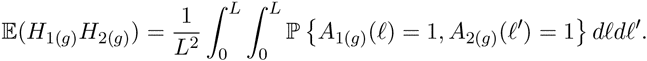

Writing these two correlation functions as

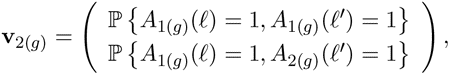

we find that

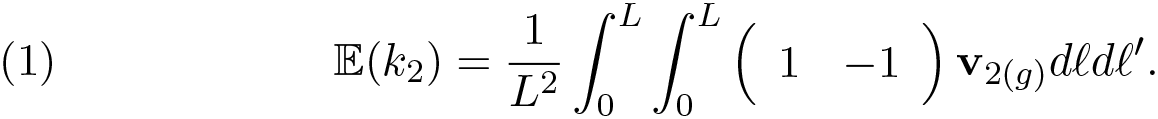

In general, the *i*^th^ *k*-statistic is an unbiased estimator of the *i*^th^ cumulant of 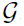, and its expectation can be written as an integral over [0, *L*]*^i^* of a linear combinations of *i*-point correlation functions. For example,

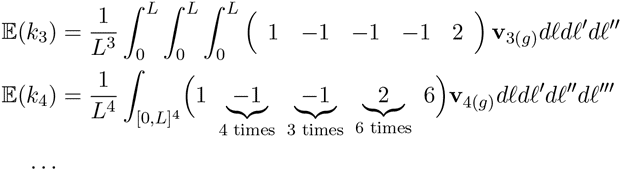

Remarkably, the linear combinations required to compute the expectations of the *k*-statistics correspond exactly to the higher-order disequilibria as defined by Bennett (1952). Furthermore, if instead the we choose to compute the expectations of the *h*-statistics, which estimate the central moments, the linear combinations would correspond to the higher-order disequilibria as defined by Slatkin (1972).

We next find the recurrence relations these correlation functions satisfy and solve them in the some special cases. In particular we will consider the case of a single admixture event *g* generations ago and the case of constant gene-flow starting *g* generations ago.

### A Single Admixture Event

We start with a simple case, where introgression only occurs in the founding generation, i.e. *s_g_* = 0 for *g* > 0. Using the random union of zygotes model, we can compute **v**_2(*g*)_ in terms of the probabilities from the previous generation:

If two sites at *ℓ* and *ℓ*′ are on the same chromosome in generation *g* + 1, then they were inherited from one chromosome from generation *g* with probability [*ℓℓ*′] and from two chromosomes from generation *g* with probability [*ℓ*|*ℓ*′]. If they are on different chromosomes, then the probability that they are descended from one chromosome in generation *g* is 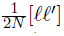 and the probability that they are descended from two chromosomes is 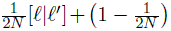 In matrix notation,

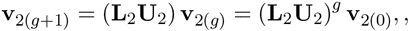

where the the recombination and drift matrices are given by

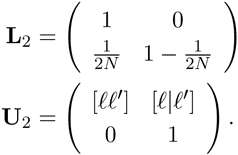

This is the the same matrix equation (Wright 1933 and Hill and Robertson 1966) derived for the decay of two-locus linkage disequilibrium. The ‘alleles’ we consider are the local ancestry at *ℓ* and *ℓ*′. To the extent possible, our notation will follow (Hill 1974), whose results for measures of multi-locus linkage disequilibria we use. The matrices **L**_2_ and **U**_2_ share (1 − 1) as a left-eigenvector, with corresponding eigenvalues 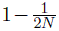 and [*ℓℓ*′]. As a result,

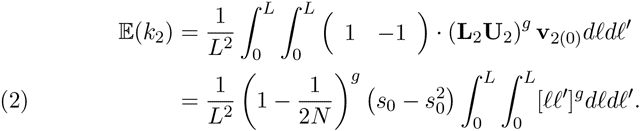

For a model using the Haldane map function, 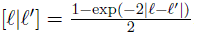, this equation becomes

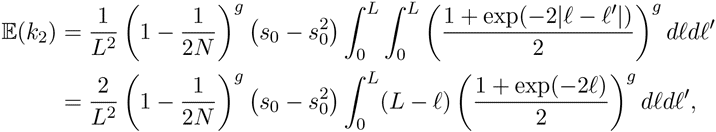

while for a model of complete crossover inteference on a chromosome of length 1 Morgan, we can get a closed form solution:

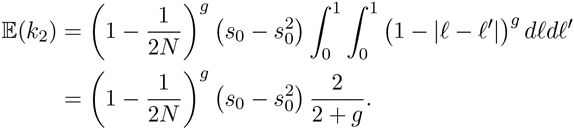

For predicting the expected sample variance, the difference between these two models is not large, as shown in figure 4. For the simulations and inference in this paper, we will ignore crossover interference, and use the Haldane map function. However, none of the mathematical results of this paper will require this assumption.

For computing higher-order correlation functions, we find a similar equation

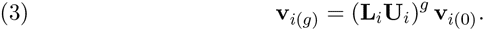

Bennett’s coefficients for higher-order linkage are left-eigenvectors of the recombination matrix **U***_i_*. For *i* = 3, it is also a left-eigenvector of the drift matrix, so we immediately get that

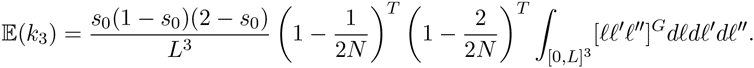

For *i* ≥ 4, this is no longer true, but the results of (Hill, 1974) can be used to compute **v***_i_*(*g*) without having to exponentiate the entire drift and recombination matrices. For example, for *k*_4_, the drift and recombination matrices are 15 × 15, but using the technique in (Hill, 1974), we only need to exponentiate a 4 × 4 matrix to compute 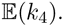.

### Varying Migration

If *s_g_* > 0 for *s* ³ 1, we obtain a modified version of Equation 3:

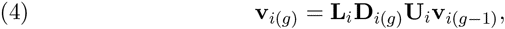

where the diagonal matrix **D***_i_*_(*g*)_ has entries giving the probabilities the set of chromosomes, *p*, in a correlation function are all from the hybrid population in the previous generation:

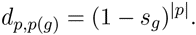

Note that if *s*_(*g*)_ is fixed, then equation (4) is linear, and can be solved using a Laplace transform.

## Inference of admixture times

The equations in the previous section can be used to develop a method of moments-estimators for admixture parameters by numerically solving the admixture parameters in terms of the expectations for the *k*-statistics. Substituting in the observed values for the *k*-statistics gives estimates for the admixture parameter(s).

However, with real data, we only have estimates of the admixture fractions, so some of the variability seen in the distribution of admixture fractions will be due to estimation variability. To account for this, we assume that the estimations errors are additive and *iid*:

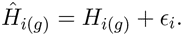

Because cumulants are additive,

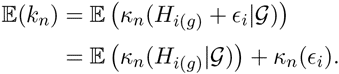

The expectations we have computed are just the term of this sum. To correct for the variability in the estimates, we need to subtract off the second term. We use a block bootstrap to estimate these effects.

One additional complication arises in dealing with genotyping data. We have assumed that we have the ancestry fractions for each haplotype in the sample, but with genotyping data, we instead have their pairwise means: 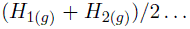. This is results in a decrease in the expectations of the *k*-statitics. Conditional on the random distribution 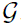, *H*_1(*g*)_, *H*_2(*g*)_, … are *iid* drawn from 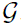. Cumulants are additive, so we use the law of total expectation to find that

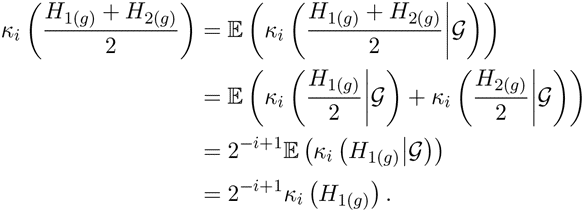

### Comparison to Verdu and Rosenberg

The recursion equations given by Verdu and Rosenberg (2011) are different from the ones we have derived. This is partly because we have accounted for the effects of genetic drift and recombination, but also because we are computing the moments of slightly different quantities.

In figure 2, we have shown the admixture fractions for five replicate populations 5, 50, and 500 generations after an admixture pulse. The variance that (Verdu and Rosenberg, 2011) compute variance over all the replicate populations, while the variance we have computed in this paper is the expectation of the variance within a single population. When *g* is small, these similar, but when *g* is large, the variance within a population goes to zero, but the variance across the replicate populations does not. This effect is shown in Figure 3. Initially, both quantities decline exponentially in *g*, but after 2*^g^* > *nLg*, the variance we predict begins to decline linearly instead. This is because variance is inversely proportional to the number of genetic ancestors of the sample. When *g* is small, the number of genetic ancestors is approximately 2*^g^*. However, the approximate number of recombination events in the sample is approximately bounded by *nLg*, so when this quantity is smaller than 2*^g^*, it provides a better approximation for the number of genetic ancestors. In this regime, the variance will decline linearly in *g*.

**Figure 2.**
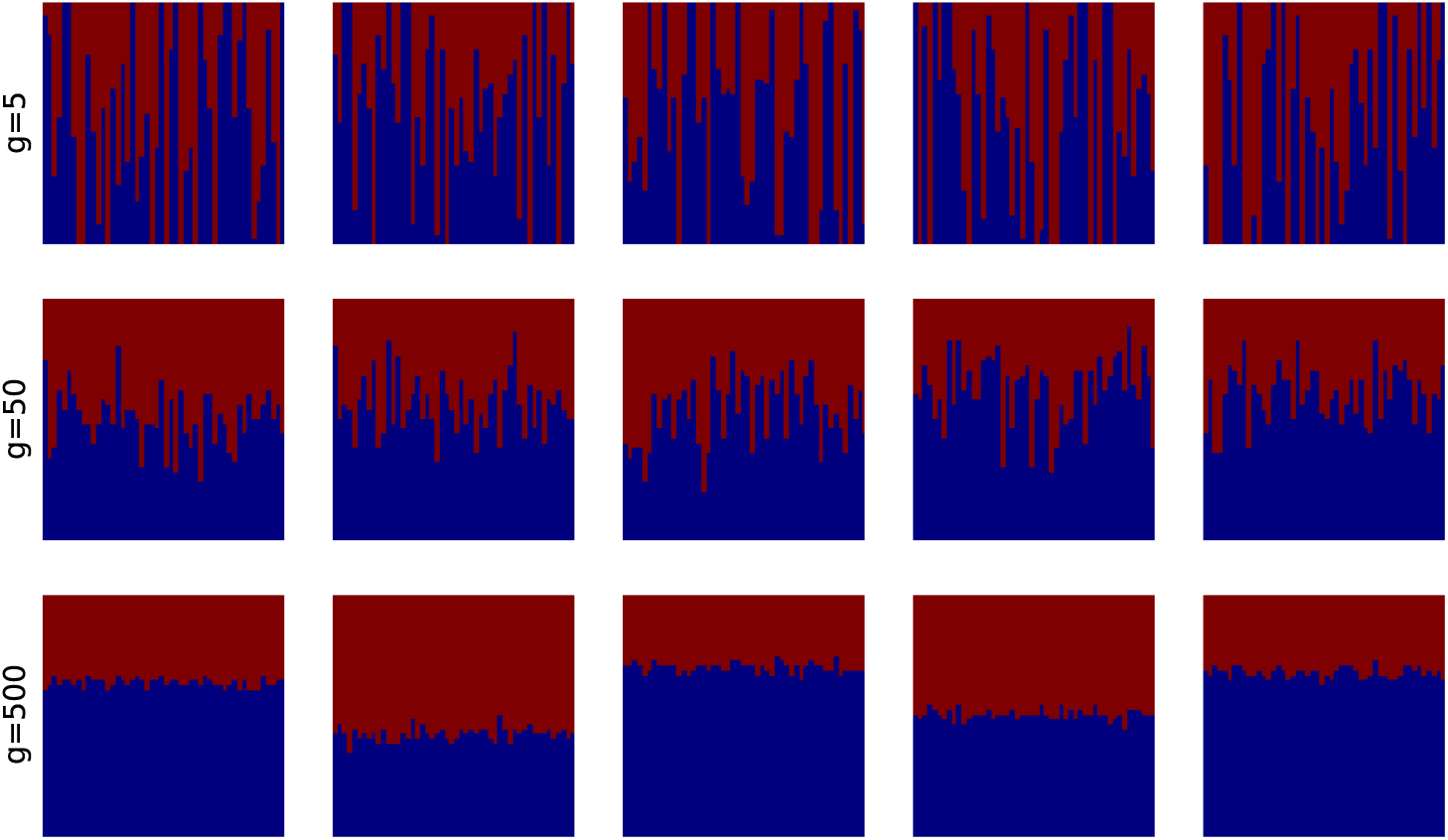
The admixture fractions of five replicate populations (each column) 5, 50, and 500 generations after an admixture pulse. As the admixture event grows more ancient, the variability within a replicate population decreases, but some variability is still maintained across the populations.

**Figure 3.**
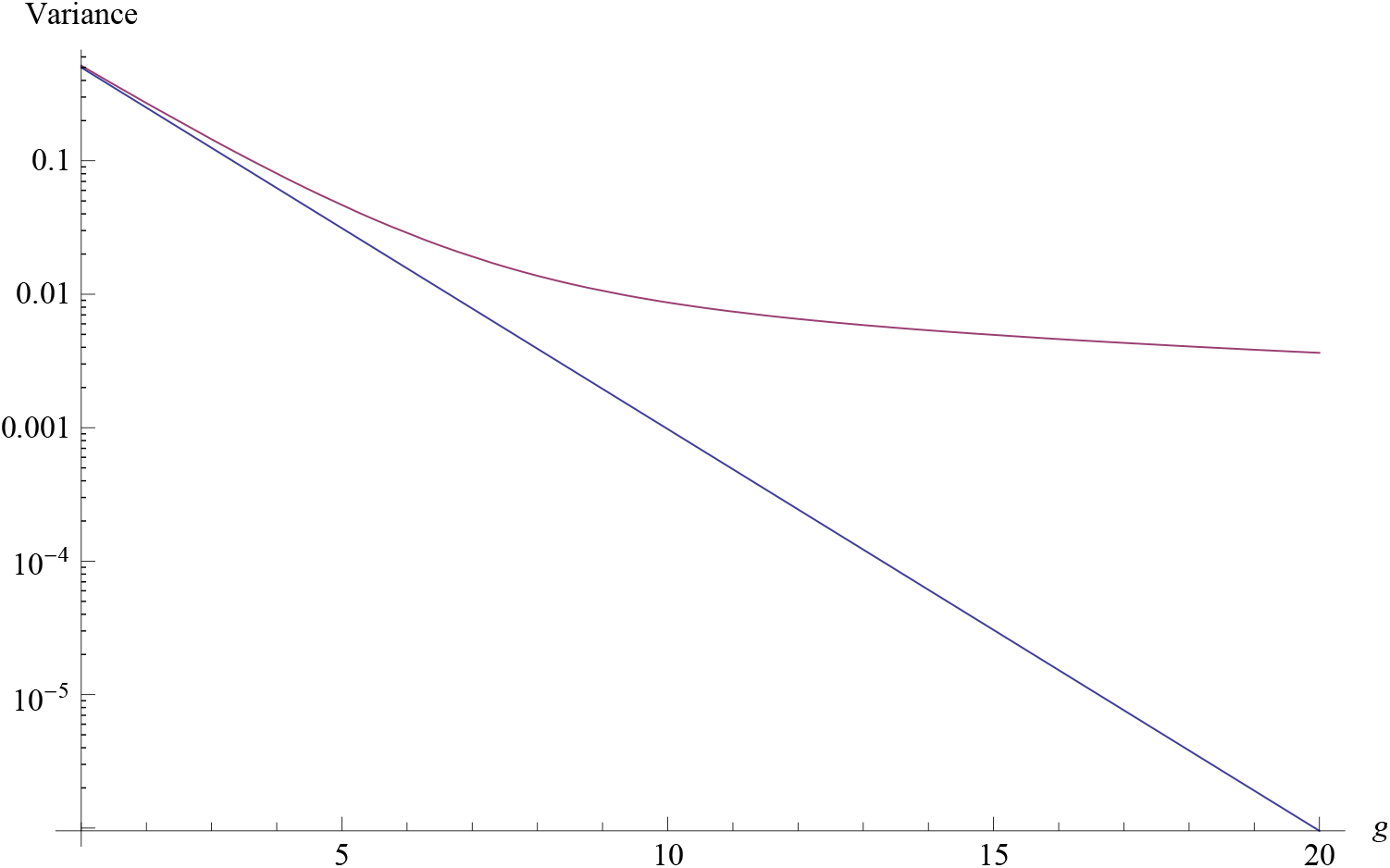
The variance predicted by Verdu and Rosenberg (2011) and equation 5, plotted on a logarithmic scale. The variance we predict (red) is always larger, but the two a very similar when *g* is small.

**Figure 4.**
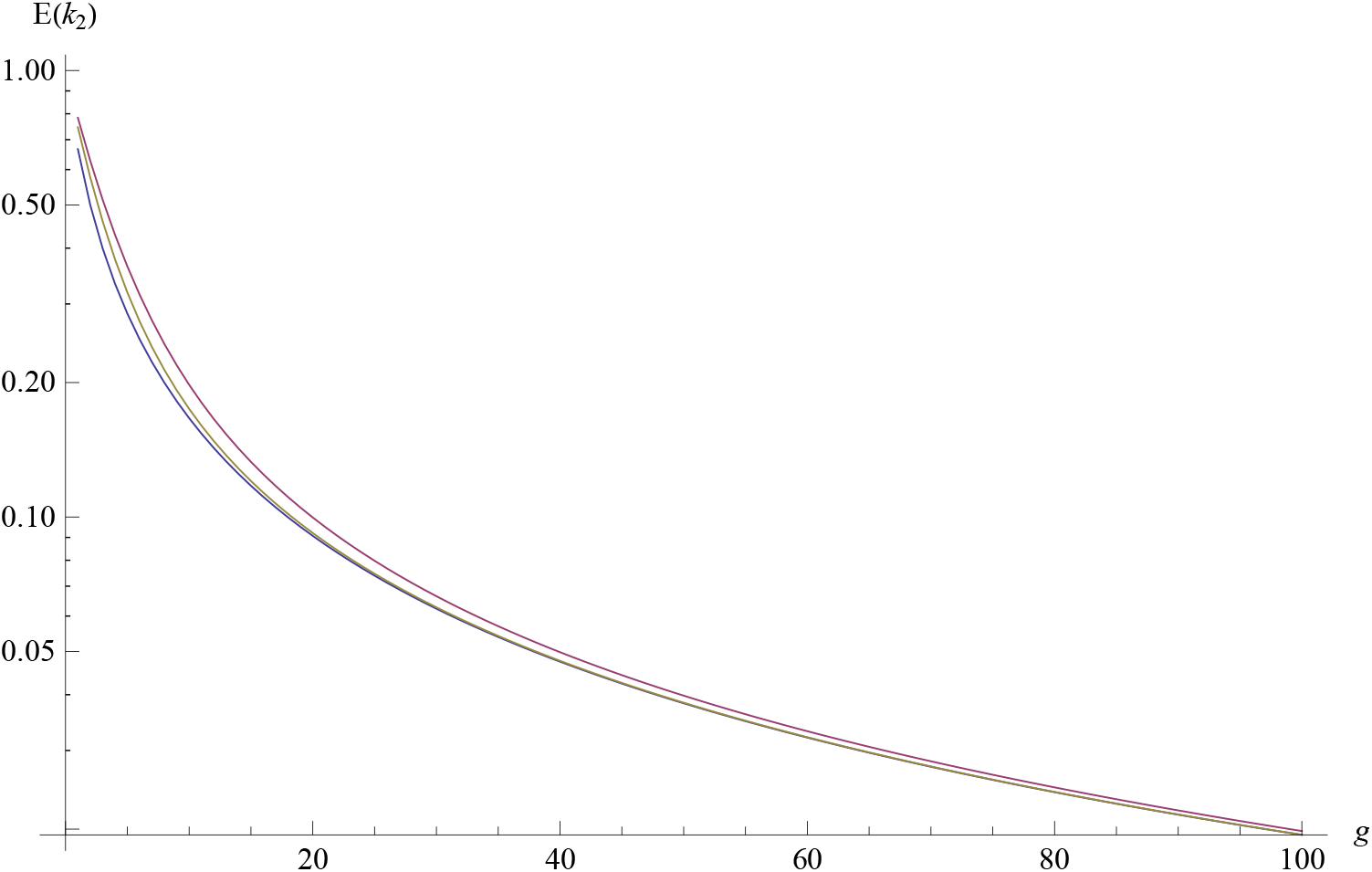
The expected sample variance given by equation 1 plotted on a logarithmic scale, for a three different map functions. We used a map distance of *L* = 1 Morgan and *N* = 10^4^. The Haldanemap function(1/2 − *e*^−2*x*^/2) is in red, the Kosambi map function (tanh (2*x*)/2) is in yellow, and the complete interence map function (*x*) is in blue. For all values of *g*, the expectations are ordered in the same order as the map functions, but the difference between the three disappears by *g* = 100.

It is also possible to compute the variance over all population replicates under our model, which allows a direct comparison to Verdu and Rosenberg (2011). In the case of one pulse of admixture, we can now solve equations 1 for 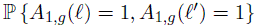 to get

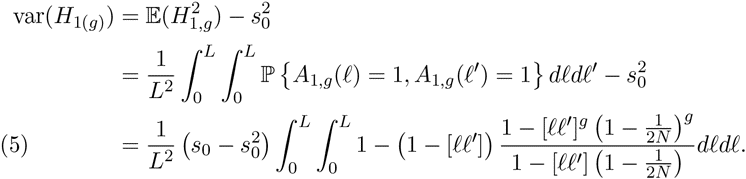

This variance and the expectation of the second *k*-statistic have the same limit as *N* → ∞, but for finite *N*, the variance is larger. This is because

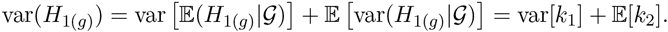

The first variance is small when *N* is large, but is always non-negative. The difference between this equation and equation 1 only becomes significant on a coalescent time scale. In the absence of genetic drift, the admixture fractions are approximately independent, becuase the samples do not share ancestors.

### Application to African American Data

We applied this method to a subset of the ASW, CEU, and YRI data from the HapMap 3 project (Consortium et al., 2010). After excluding children from trios, there were the genotypes for 49 ASW,113YRI, and 112CEU individuals. We estimated the admixture fractions using the supervised learning mode of Admixture, with the CEU and YRI individuals assigned to separate clusters. The sampling distribution of the admixture fractions was estimated using the block bootstrap with 10^4^ replicates and 2678 blocks, giving a block size of approximately 10 CM. The admixture fractions for the 49 ASW samples are shown in Figure 1 and the observed *k*-statistics are given in table 6.

We assumed a 3-parameter model of constant admixture. For *g_start_* ≤ *g* ≤ *g_stop_*, *s_g_* = *s* with *s_g_* = 0 elsewhere. By matching the block-bootstrap corrected *k*_2_ and *k*_3_ to the predictions of equation 1, we obtained a point estimates of

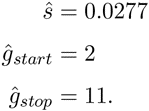

We obtained confidence intervals, shown in Figure 5, by simulation. For each cell in the grid, we simulated 10^3^ replicates under the corresponding *g_start_* and *g_stop_*, with 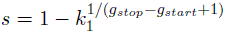. For each replicate, we computed the *k*_2_, *k*_3_, and *k*_4_ statistics. A cell was then included in the confidence interval if and only if the corrected *k*_2_, *k*_3_, and *k*_4_ statistics from the HapMap data fall inside a centered interval containing 98.7% of the probability mass of the simulated distribution. This mass was chosen so that under the Bonferroni correction for three tests, there is at least a 95% chance of including the true parameter values in the confidence region.

**Figure 5.**
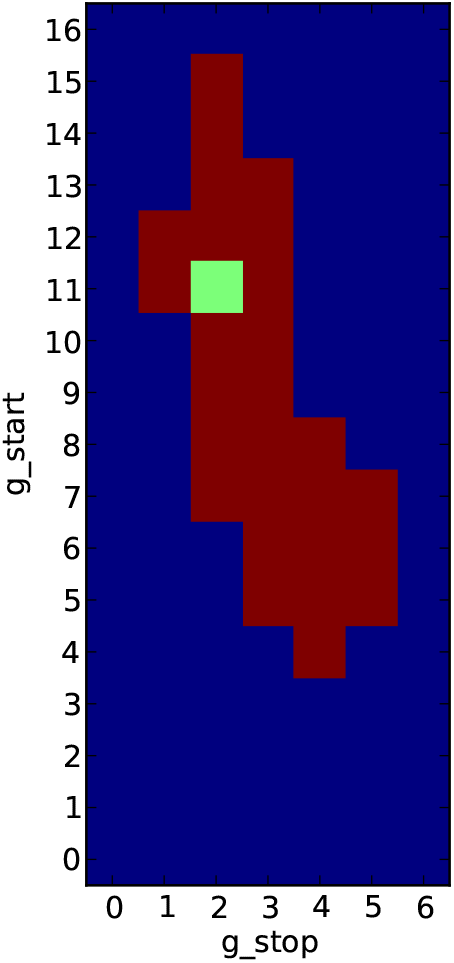
95% confidence region for a model with constant admixture from generations *g_start_* to *g_stop_*. The point estimate of *g_start_* = 11 and *g_stop_* = 2 generations ago is colored green.

**Figure 6.**
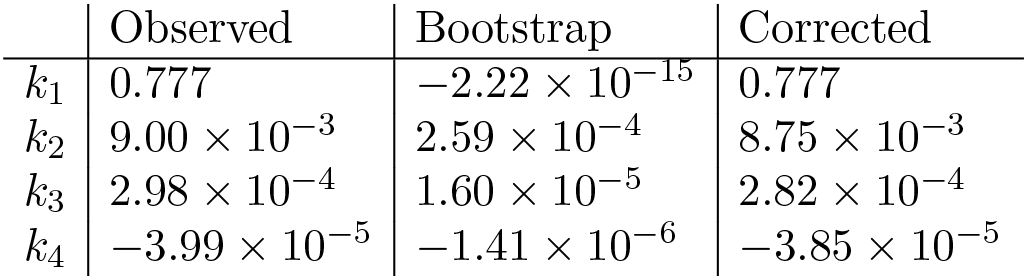
*k*-statistics

The point estimates for *g_start_* and *g_stop_* correspond to the values for which the observed *k*-statistics are closest to their simulated medians.

## Discussion

We have extended the mechanistic model of Verdu and Rosenberg (2011) to account for recombination and genetic drift. Doing so allows us to apply the predictions of this model to data. This mechanistic model allows for a large number of parameters. For the purposes of inference, it seems that imposing constraints, i.e. a small number of pulses or constant admixture, will be needed to narrow the search space.

In this paper, we have assumed that admixture only comes from one source population, this need not be the case. To account for admixture from multiple source populations, equation 1 must be modified to account for the probability that haplotypes trace their descent to multiple source populations. Algorithmically, this is feasible, but the notation is cumbersome. The resulting equations are given in the appendix, along with the equations for computing expectations of higher-order *k*-statistics.

Applications of the method to African-American HapMap data provides estimates of the time since admixture between people of Europe and and African descent in America. Notice that the confidence set for the admixture parameters does not include values of *g_stop_* = 0. We interpret this as evidence that admixture rates have declined the last few generations. The point estimate of time gene-flow stopped is *g_stop_* = 2. This probably reflects a more gradual reduction in gene-flow within the last 5 generations or so, rather than a discrete stop in gene-flow 2 generations ago. The discreteness is enforced by the model. Also notice that admixture before 15 generations ago can be rejected. With a generation time of 25-30 years, this corresponds to 325-400 years, and is in good accordance with the historical record. The point estimate of the time of first admixture is 11 generations, or approx. 275-330 years ago.

Structure analyses have become one of the most commonly applied tools in population genomic analyses. The theory developed in this paper allows users of structure analyses to interpret their data in the context of a model of admixture between populations, and should find use in many studies aimed at understanding the history of populations.

## Appendix

These are the matrices for computing 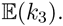. The matrices for computing 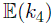 are 15 × 15 and not given here, but can be found in (Hill, 1974).

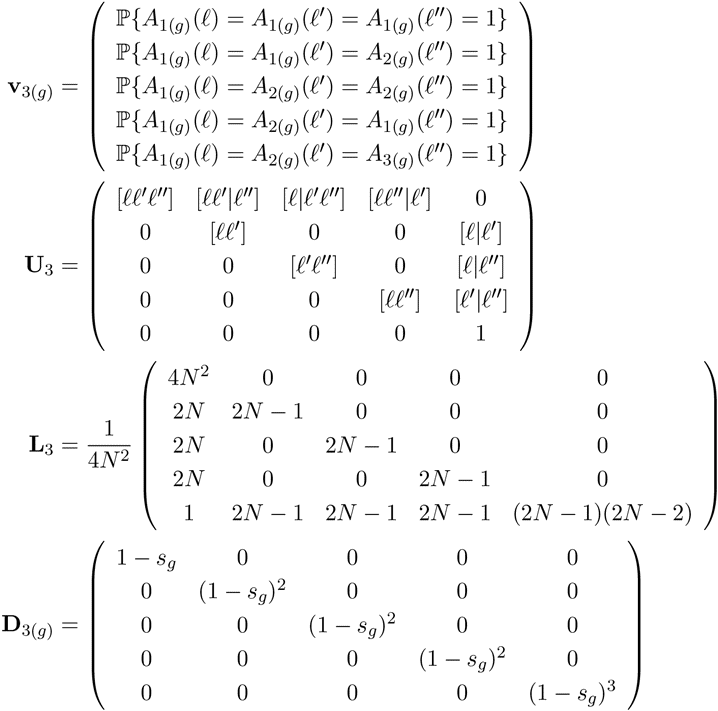

When there is migration from both source populations, the recursion relations for the *i*-point correlation functions will depend on *i* − 1-point, *i* − 2-point, … correlations functions as well. As as example, consider the case of **v**_2(*g*)_. Let the introgression probability from the second source population be given by *t_g_*. The recursion equation for **v**_2(*g*)_ now also depends on **v**_1(*g*)_.

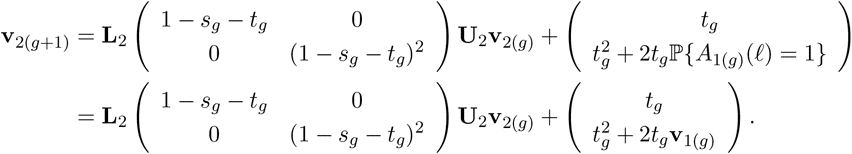

Similarly, the recursion equation for **v**_3(*g*)_ depends on **v**_2(*g*)_ and **v**_1(*g*)_.

## References

David H Alexander, John Novembre, and Kenneth Lange. Fast model-based estimation of ancestry in unrelated individuals. Genome Research, 19(9): 1655–1664, 2009.

John Bennett. On the theory of random mating. Annals of Eugenics, 17(1): 311–317, 1952.

International HapMap 3 Consortium et al. Integrating common and rare genetic variation in diverse human populations. Nature, 467(7311):52–58, 2010.

Daniel Falush, Matthew Stephens, and Jonathan K Pritchard. Inference of population structure using multilocus genotype data: linked loci and correlated allele frequencies. Genetics, 164(4):1567–1587, 2003.

Simon Gravel. Population genetics models of local ancestry. Genetics, 191 (2):607–619, 2012.

William G Hill. Disequilibrium among several linked neutral genes in finite population i. mean changes in disequilibrium. Theoretical Population Biology, 5(3):366–392, 1974.

Mason Liang and Rasmus Nielsen. The lengths of admixture tracts. Genetics, pages genetics–114, 2014.

Brian K Maples, Simon Gravel, Eimear E Kenny, and Carlos D Bustamante. Rfmix: A discriminative modeling approach for rapid and robust localancestry inference. The American Journal of Human Genetics, 93(2): 278–288, 2013.

Marilyn Menotti-Raymond, Victor A David, Solveig M Pflueger, Kerstin Lindblad-Toh, Claire M Wade, Stephen J OBrien, and Warren E Johnson. Patterns of molecular genetic variation among cat breeds. Genomics, 91 (1):1–11, 2008.

John E Pool and Rasmus Nielsen. Inference of historical changes in migration rate from the lengths of migrant tracts. Genetics, 181(2):711–719, 2009.

Alkes L Price, Nick J Patterson, Robert M Plenge, Michael E Weinblatt, Nancy A Shadick, and David Reich. Principal components analysis corrects for stratification in genome-wide association studies. Nature Genetics, 38(8):904–909, 2006.

Jonathan K Pritchard, Matthew Stephens, and Peter Donnelly. Inference of population structure using multilocus genotype data. Genetics, 155(2): 945–959, 2000.

Noah A Rosenberg, Jonathan K Pritchard, James L Weber, Howard M Cann, Kenneth K Kidd, Lev A Zhivotovsky, and Marcus W Feldman. Genetic structure of human populations. Science, 298(5602):2381–2385, 2002.

Montgomery Slatkin. On treating the chromosome as the unit of selection. Genetics, 72(1):157–168, 1972.

Hua Tang, Jie Peng, Pei Wang, and Neil J Risch. Estimation of individual admixture: analytical and study design considerations. Genetic epidemiology, 28(4):289–301, 2005.

Hua Tang, Marc Coram, Pei Wang, Xiaofeng Zhu, and Neil Risch. Reconstructing genetic ancestry blocks in admixed individuals. The American Journal of Human Genetics, 79(1):1–12, 2006.

Paul Verdu and Noah A Rosenberg. A general mechanistic model for admixture histories of hybrid populations. Genetics, 189(4):1413–1426, 2011.

Baowei Zhang, Ming Li, Zejun Zhang, Benoît Goossens, Lifeng Zhu, Shanning Zhang, Jinchu Hu, Michael W Bruford, and Fuwen Wei. Genetic viability and population history of the giant panda, putting an end to the evolutionary dead end? Molecular biology and evolution, 24(8):1801–1810, 2007.

